# Strong negative reinforcement interferes with visual learning in a solitary pollinator

**DOI:** 10.1101/2025.06.18.660396

**Authors:** Anupama Nayak Manel, James Jonathan Foster, Anna Stöckl

**Affiliations:** Department of Biology, University of Konstanz, Konstanz, Germany; International Max Planck Research School for Quantitative Behaviour, Ecology and Evolution (IMPRS-QBEE), Max Planck Institute of Animal Behaviour, Radolfzell, Germany; Centre for the Advanced Study of Collective Behaviour, University of Konstanz, Konstanz, Germany; Zukunftskolleg, University of Konstanz, Konstanz, Germany

## Abstract

Visual learning in insects can be strongly influenced by the underlying context and conditions. Improved colour learning has been demonstrated with differential conditioning using reward-aversion paradigms in eusocial insects like wasps, bumblebees, and honeybees, where substances that induce strong aversion, such as quinine, enable more accurate learning. Although the associations created during learning are intrinsically linked to the reinforcements used, few studies have compared the role of negative reinforcements in shaping these associations, and their relevance across insect groups with different life histories.

In our study, we compared the effects of aversive substances for learning in the hummingbird hawkmoth (*Macroglossum stellatarum)*, a solitary pollinator that relies on vision for foraging. Linking to previous learning studies in insects, we combined a sugar-rewarded target with a distractor that was paired with either quinine, salt, citric acid, water, or was presented with no aversive substance. Learning was assessed by conditioning hawkmoths to invert a strong colour preference between perceptually close or distant colour pairs, to provide tasks with disparate challenges. Contrary to results from eusocial insect species, hawkmoths trained with quinine were worse at switching preferences between similar colours compared to training with citric acid or appetitive-only differential conditioning. Learning success was associated with the animals’ foraging plasticity, where negative reinforcements were found to suppress exploration during foraging. Furthermore, we show that quinine interfered with sucrose perception, potentially impairing target acquisition during conditioning. Our results provide insights into the impact of negative reinforcements on solitary foragers and highlight the role of sensory ecology and life history in shaping learning outcomes.

## INTRODUCTION

Visual learning in insects can be highly plastic, with behavioural outcomes varying in response to experiential details such as learning durations (Stach & Giurfa, 2005), or the context in which learning takes place (Avarguès-Weber & Giurfa, 2014; Buatois et al., 2017; Dyer & Chittka, 2004; Dyer & Howard, 2023; Giurfa, 2004; Howard et al., 2019; Kelber, 2010a; Matsumoto & Mizunami, 2002). In particular, the conditions in which stimuli are presented are known to alter the speed and accuracy of learning depending on whether the animal experiences absolute conditioning to a single target, or differential conditioning in the presence of a rewarded target stimulus along with a distractor stimulus associated with a neutral or negative feedback (Avarguès-Weber & Giurfa, 2014). Differential conditioning enables narrower colour discrimination and pattern recognition where absolute conditioning is less effective (Dyer & Chittka, 2004; Giurfa, 2004). Such improved learning performance has been attributed to the reward and punishment dynamic of differential conditioning, which could motivate animals to be more attentive to the task due to the adverse consequences of incorrect choices (Avarguès-Weber & Giurfa, 2014). Moreover, differential learning provides a richer set of information to enable selective association, since both, the positively associated target and negatively associated distractor stimuli can be learnt and distinguished simultaneously.

Since the strength of association with the target and distractor is a key aspect of differential learning, the reinforcement type paired with the distractor plays a critical role in the conditioning process. Across insect groups, various substances, such as water, salt solution, or bitter alkaloids (Hymenoptera: Josens et al., 2009; Lacombrade et al., 2023; Orthoptera: Matsumoto, 2022; Lepidoptera: Rodrigues, 2016; Young et al., 2024) have been used as negative reinforcements. Alternatively, appetitive-only conditioning, where the target is rewarded while the distractor is not associated with any substance, has also been used for insect training (Goyret et al., 2008; Kelber, 2002). Previously, in eusocial Hymenoptera, colour discrimination between perceptually similar– and thus harder to learn– stimuli was shown to improve by using the aversive alkaloid quinine as a negative reinforcer, compared to using water (Avarguès-Weber et al., 2010; Rodríguez-Gironés et al., 2013), or with no negative reinforcement (Dyer & Howard, 2023). However, even within Hymenoptera, the use of quinine as a negative reinforcement remains contested since aside from being toxic to the insects (Ayestaran et al., 2010), its efficacy is also context-dependent: although it improved learning in untethered honeybees, it had no effect on learning in harnessed bees, which sometimes even ingested the toxic bitter compound (Ayestaran et al., 2010; de Brito Sanchez et al., 2015).

The wide disparity of learning conditions, in addition to the limited evidence for assessing their suitability across insect models, makes it a challenge to interpret learning outcomes across species. To understand which learning strategies are shared across insects, and which are specific adaptations to a species’ lifestyle and environment, it becomes crucial to understand how reinforcing substances might present disparate qualities for species with different sensory systems and life histories. For instance, phytophagous insects specialised on certain host plants may have different tolerances for specific secondary compounds produced by the plant. The fly, *Drosophila sechellia*, a specialist adapted to the toxic fruits of *Morinda citrifolia*, shows reduced aversion to the bitter compounds rejected by other Drosophilidae (Reisenman et al., 2023). Similarly, frugivorous butterflies were shown to be more receptive to amino acids in their diets than nectarivorous butterflies (Ravenscraft & Boggs, 2016). Such differences, even between closely related species, question whether there is a ubiquitous role of appetitive and aversive substances in insect learning.

To assess how different negative reinforcements influence visual discrimination learning in a solitary insect, we differentially conditioned the day-active hummingbird hawkmoth, *Macroglossum stellatarum (Fig. 1)*. *M. stellatarum* are generalist nectar feeders that rely on vision for locating and interacting with flowers (Balkenius, Rosén, et al., 2006; Kelber, 1997; A. Stöckl et al., 2016). They represent a large group of solitary insects that occupy a key pollinator niche (Johnson et al., 2017; Singh et al., 2025), sharing many habitats and floral hosts with eusocial pollinators. Both behavioural and neural correlates of visual foraging in this species have been extensively studied (Balkenius, Kelber, et al., 2006; Kannegieser et al., 2024; Kelber, 2002; A. Stöckl et al., 2016; A. L. Stöckl et al., 2017). Previously, they have been found to make rapid visual associations via absolute conditioning. However, their learning outcomes were limited by their strong preferences for certain pattern and colour features (Kelber, 2002). To test whether conditioning with negative reinforcements could improve their visual association learning, we differentially conditioned the hawkmoths using substances in the taste categories-sour, salt, bitter, and water (neutral) – as negative reinforcements. These negative reinforcements were compared with an appetitive-only condition. Hawkmoths were conditioned to invert a strong colour preference in two visual discrimination tasks: one with perceptually close and another with perceptually distant colours. With this preference inversion paradigm, we addressed whether the type of negative reinforcement altered learning outcomes, and how the effects of reinforcement varied with the challenge of the visual discrimination task.

**Fig. 1.**
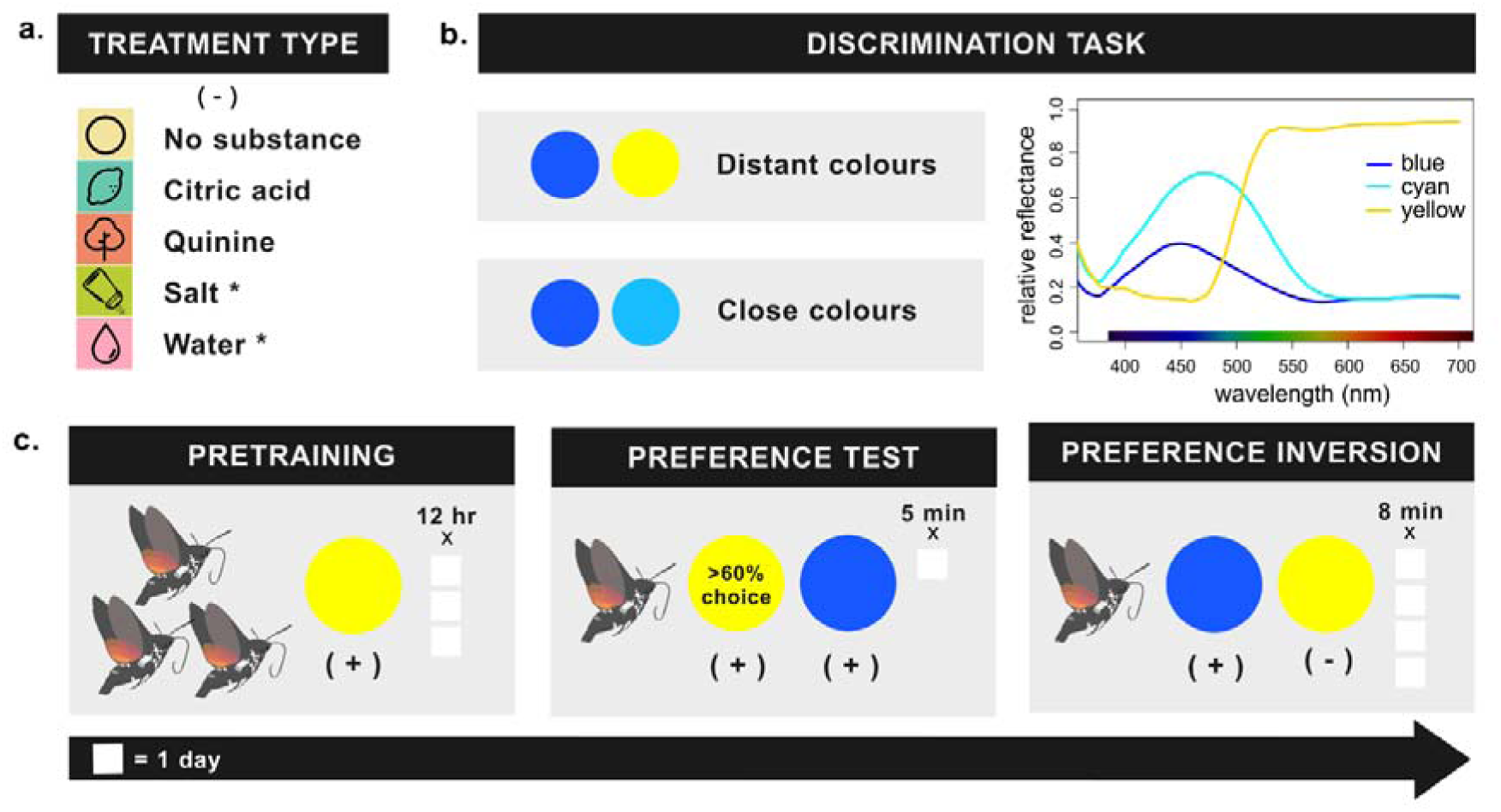
Comparing negative reinforcement types for colour discrimination learning in the diurnal hawkmoth, *Macroglossum stellatarum*. **a**. Learning was compared across differential conditioning protocols with various treatment types (-) using aversive substances common in insect training, as well as an appetitive-only condition with no aversive substance. Asterisks indicate treatments used only in the distant colour discrimination task. **b.** Colour preference inversion was compared across two discrimination tasks with stimuli that were perceptually distant (blue-yellow) and close (blue-cyan). Relative reflectance of the colours used in the experiment (see Methods for details on measurements, and Fig. S1 for distances modelled in the hawkmoths’ colour space). **c.** The experimental paradigm was divided into 3 phases: moths were pretrained to artificial flowers of a single colour in groups and then individually tested for a preferred colour in a dual choice assay. Following this, their preferred colour was paired with a single treatment type (-) as a negative reinforcer and the unpreferred colour was rewarded with sugar (+). The figure depicts the distant colour discrimination task as an example, with yellow as the moth’s preferred colour. Durations for each phase are depicted as white boxes representing a single day.

## Materials and Methods

### Animals

Hummingbird hawkmoth (*Macroglossum stellatarum*) populations were maintained indoors at the University of Konstanz, Germany at 23 °C and 40-60% humidity with a 14:10 h light:dark cycle. Populations were supplemented with wild-caught moths when available. Caterpillars were raised on their native host plant *Gallium* sp.. Adults were fed in flight cages (60 cm × 60 cm × 60 cm) with 20% sucrose-water solution from gravity feeders. Multiple feeders in blue and yellow colours were provided simultaneously to discourage colour associations.

Animals used in the experiment were 3-6 weeks of age. Moths were placed in plastic vials (8 cm x 3.5 cm length x diameter) and kept in the dark when not participating in the experiments. The vials were marked to track individual identity across the experiment.

### Experimental Setup

Experiments were performed in a flight cage of 60 cm × 60 cm × 60 cm dimension at 23 °C. The cage was lit from above with four 60 cm long daylight-like fluorescent tubes (Osram L 18 W/965 Biolux Tageslicht G13). Lights were connected to an electrical ballast (GloMat 2 × 40 W, Hagen), to increase the flicker frequency of the fluorescent tubes above 25 kHz, beyond the resolvable range of hawkmoths (Stöckl et al., 2019). A USB camera fixed to the upper corner of the cage was used to record the experiments. It was controlled by the ContaCam software (ContaCam v9.0.9beta4, https://www.contaware.com) to record videos at 30 fps.

Artificial flowers were constructed by pasting coloured discs (38 mm diameter) on cylindrical plastic pedestals (8 cm x 3.5 cm) wrapped with unbleached paper. Discs in Blue (CMYK 91:66:0:0), Yellow (CMYK 0:0:100:0), and Cyan (CMYK 91:26:0:0) were designed in the vector graphics software Inkscape (Inkscape v1.3 2023, https://inkscape.org). The colours were laser printed (imageRUNNER ADVANCE DX C3926i, Canon) on unbleached paper (‘Classic White’, Steinbeis) and laminated with matte foil (S-PP525-22 matt, Peach, PRT GmbH). To provide optic flow during flight, matte-laminated dead-leaves patterns (Lee et al., 2001) were used on the floor of the cage. During the test and conditioning phase of the experiments, the flowers were placed on a dead-leaves pattern covered cardboard platform (30 cm × 50 cm × 50 cm) higher off the ground of the cage to decrease the flight height and thereby get increased feeding responses.

Irradiance measurements of the flower colours, background, and lighting were taken using a calibrated Miniature Spectrometer (OceanHDX, Ocean Optics Inc., Dunedin, FL, USA) connected to a UV-VIS optical fibre (P400-2-UV-Vis). Relative reflectance of the stimuli was then calculated with respect to a 99% white standard (Spectralon diffuse reflectance standards, Labspehere Inc., USA). All spectral measurements were processed and visualized in RStudio (Rstudio Team, v2023.09.1) using the pavo 2.2.0 package (Maia et al. 2019) (*Fig. S1a*).

To map the colour distances, quantum catches for the photoreceptors were calculated by integrating the sensitivity peaks of *M. stellatarum* at 349, 440, and 521 nm (Telles et al., 2014), the cage illumination, and irradiance measures of the stimuli (Renoult et al., 2017). The ‘sensmodel’ function from the pavo package was used to set the sensitivity values for the hawkmoths, which were then input into the ‘vismodel’ function to calculate photoreceptor stimulation based on von-Kries colour corrections (Maia et al., 2019). Relative chromatic and achromatic distance between the colours were calculated as noise-weighted Euclidean distances measured in Just Noticeable Differences (JNDs) (Table S1, *Fig. S1b*), which is a contrast threshold that uses the RNL model (Vorobyev & Osorio, 1998). Relative photoreceptor densities were assigned as 1:1:7 for the small, medium, and long wavelength receptors (Kelber et al., 2003).

### Experimental Procedure

Hawkmoths were differentially conditioned to invert a strong colour preference using various treatment types as the negative reinforcement. By using quinine hydrochloride (120 mM), citric acid (1.04 M), and sodium chloride (3 M) solution, distilled water, and no substance (appetitive-only condition), we combined treatments from insect literature (Avarguès-Weber et al., 2010; Kelber, 2002) to compare a variety of negative reinforcements. We assessed learning in 2 visual discrimination tasks associated with disparate learning challenges (Dyer & Chittka, 2004; Giurfa, 2004). For the easier task we used blue and yellow which are further apart in hawkmoths’ colour space (distant colours), and for the harder discrimination task we used blue and cyan (close colours) (*Fig. 1b, see Fig. S1b for colour spaces*) as the conditioned stimuli (CS). Individual hawkmoths experienced a single treatment and discrimination type with the full paradigm divided into 3 phases – Pretraining, Preference test, and Preference inversion *(Fig. 1c)*.

In all but the pretraining phase, animals were tested individually in the flight cages. Food (20% sucrose solution) was only available during the trials and for the remainder of time, animals were kept in vials under dark conditions. Artificial flowers within the flight cage were randomly arranged at least 5 cm away from each other, and their positions were changed within and between trials to avoid place learning. The negative reinforcement substances and sugar rewards were pipetted out at several points on the surface of the flower as small droplets (20-30 µl), requiring the moths to make multiple choices before satiation. The flower surfaces were cleaned between all trials. A trial commenced when the moth began its flight, and during the trial every new approach to an artificial flower that ended with a proboscis contact to the surface was scored as a choice. Repeated contacts on the same surface while hovering or repositioning around the flower were not included as additional choices. All trials were video documented.

#### Pretraining

To habituate the moths to the artificial flowers, groups of 5-8 moths were pretrained to the setup by sequentially switching their regular gravity feeders with artificial flowers across 3 days. On the first day the moths had one feeder and 2 artificial flowers at different elevations, and by the third day only 3 artificial flowers were available to feed from. Each group was pretrained to flowers of a single randomly chosen colour. Either yellow or blue was used for the distant colour experiments, and either blue or cyan were used for the close colour experiments. During this phase, the moths could feed ad libitum on 20% sucrose from the surface of the flowers for 10-12 hours each day. To ensure the flowers always contained sucrose during visits, they were removed from the cage in the evenings. On the evening of the third day, moths were packed into individual vials and kept in a dark box for 48 hours to motivate feeding during the next phase.

#### Preference test

Moths were tested for a preferred feeding colour between the two colours assigned to their discrimination type, blue and yellow (distant colours), or blue and cyan (close colours). The identity and sex of the moth were noted, along with the time of start of each experiment. Six artificial flowers, 3 of each colour, were arranged in the flight cage with 20% sucrose drops available on all the flowers. All choices were rewarded to prevent the animals from forming negative associations with their preferred colour, so the animals could feed during the trial rather than in a non-experimental setting. Flower choices were scored by the experimenter across 5 minutes of flight. Only animals with a relative colour preference, i.e. at least 60% of choices on one of the colours, were used in the following preference inversion phase.

#### Preference inversion

Hawkmoths were differentially conditioned to their unpreferred colour as determined by the preference test. For the distant colour experiments, the artificial flowers with the unpreferred colour (CS+) were paired with a reward of 20% sucrose while the preferred colour (CS-) was associated either with citric acid (n=18), quinine hydrochloride solution (n=16), sodium chloride solution (n=18), distilled water (n=14), or, in the case of the appetitive-only conditioning, no substance (n=14).

For the close colour (blue-cyan) discrimination condition, only citric acid (n=13), quinine (n=15), and no substance (n=14) were used, as these treatment types emerged as most relevant from the distant colour experiments.

The moths were differentially conditioned in 4 trials, one per day, during which they had access to the artificial flowers (3 x CS+ and 3 x CS-) for 8 minutes of flight. This duration was determined to be a suitable interval for most moths to approach the flowers and make sufficient choices until satiation based on pilot experiments. Only sessions in which moths made at least one choice were included as trial days.

#### Control experiment

For the appetitive-only conditioning, only the CS+ was presented with droplets. To assess whether the animals associated the droplets rather than the colour as a rewarding cue, we altered the preference inversion phase with a test on the fourth day where all the flower surfaces were bare *(Fig. S2a)*. For this test, choices were recorded for 5 minutes of flight, rather than 8 minutes, to account for changes in preference associated with finding no rewards on the CS+. This control was conducted for the appetitive-only condition (n=9) with the close colour setup where the results showed a significant relation between learning and treatment type.

### Data Analysis

Data was analysed separately for the distant colour and close colour discrimination experiments. Moths that participated for at least 3 days of differential conditioning were included in the analysis. All statistical analyses were conducted in R (R Core Team, 2021), and plots were created with the ggplot2 package (Wickham, 2011).

To compare the number of individuals that succeeded in the preference inversion task between treatments, we assigned a success criterion to moths that shifted their colour preference in relation to the 60% preference criterion in the preliminary test, i.e., individuals with greater than 40% CS+ choices (the initially non-preferred colour) on the fourth day. A pairwise Fisher’s exact test with Holm corrections for multiple testing was then implemented to compare the numbers of successful and unsuccessful moths between treatments *(Fig. S3b)*.

#### Choice rates

Generalized linear-mixed models (GLMM) fitted with the ‘glmer’ function from the lme4 package were used to test if the correct choice rate varied significantly between treatment types, and across the differential conditioning trials. Choice data from the differential conditioning was fitted to a binomial family with treatment type, day, preferred colour, sex, and the choice rate from the preference test as fixed effects, and individual ID as the random effect. Residuals from the model were simulated using the ‘simulateResiduals’ function from the DHARMa package (Hartig, 2018) to validate model accuracy. Stepwise reduction of the full model was conducted with likelihood ratio tests (LRT) via the ‘drop1’ (Crawley, 2002) function, and the random effect structure of the final model was selected based on Akaike information criterion (AIC) comparisons. The simplest adequate model was formulated as

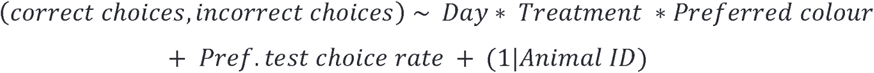

Preferred colour as a predictor was removed from the close colour experiment model where only blue was the CS-. The ‘Anova’ function from the car package (Fox and Weisberg, 2019) with Wald’s Chi-square test was used to determine the significance of the fixed effect variables on the choices. Post-hoc comparisons were then conducted to identify pairwise differences using a Sidak (Šidák, 1967) correction for multiple comparisons with the emmeans package (Lenth et al. 2019) *(Fig. 2,3)*.

**Fig. 2.**
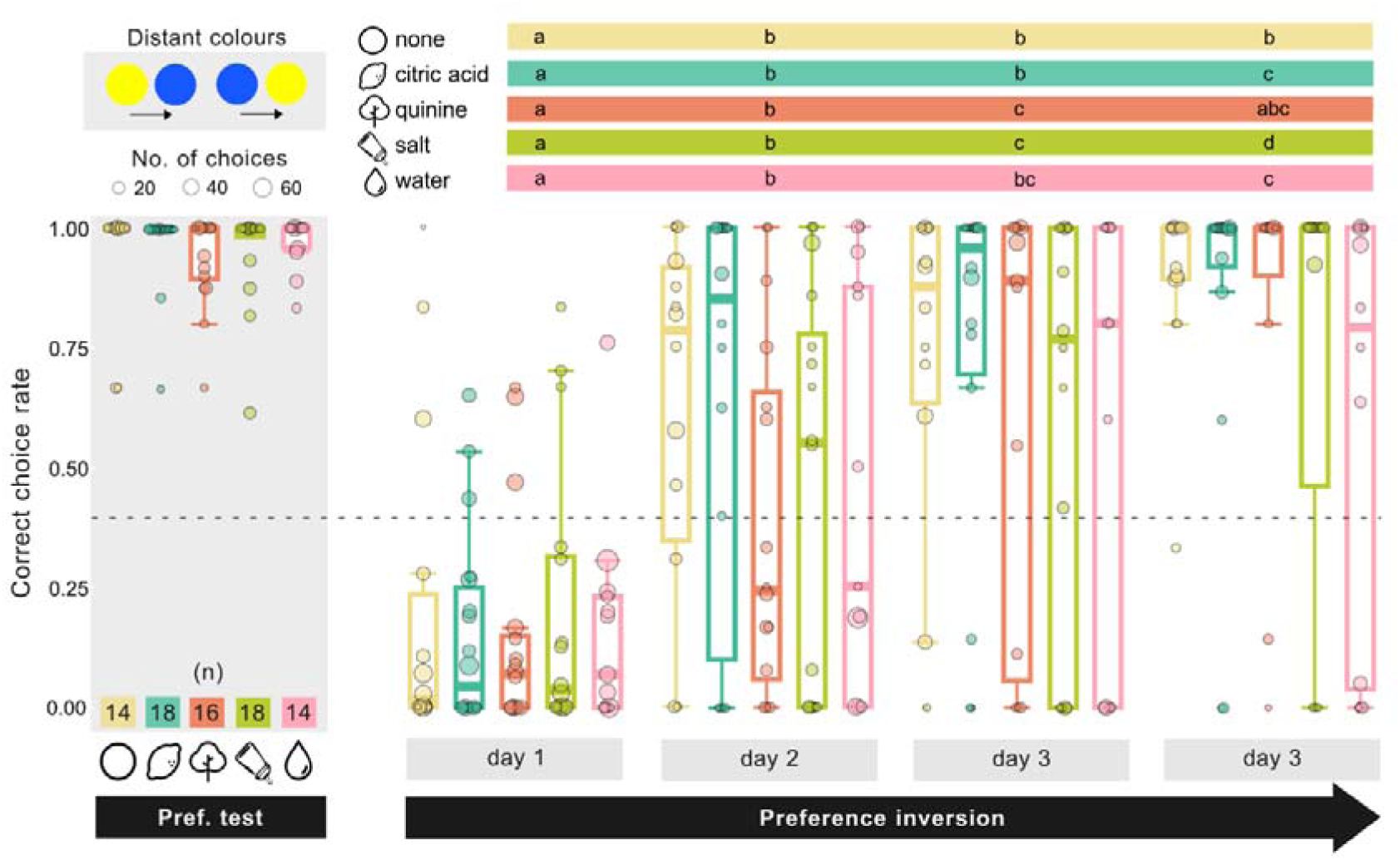
Preference inversion in the distant colour experiments. Correct choice rates represent feeding on the preferred colour during the preference test (grey highlight), and on the unpreferred colour (CS+) during the preference inversion phase. Choice rates of individuals are denoted by circles with the size representing the number of choices made in a day (see legend). Statistical results (GLMM, Fig. S4a) of differences in choices rates between days within a treatment type are indicated on the treatment-specific coloured bars where days with no differences *(p>0.05)* share the same letter. Pairwise comparisons of treatments within a day were not significant.

**Fig. 3.**
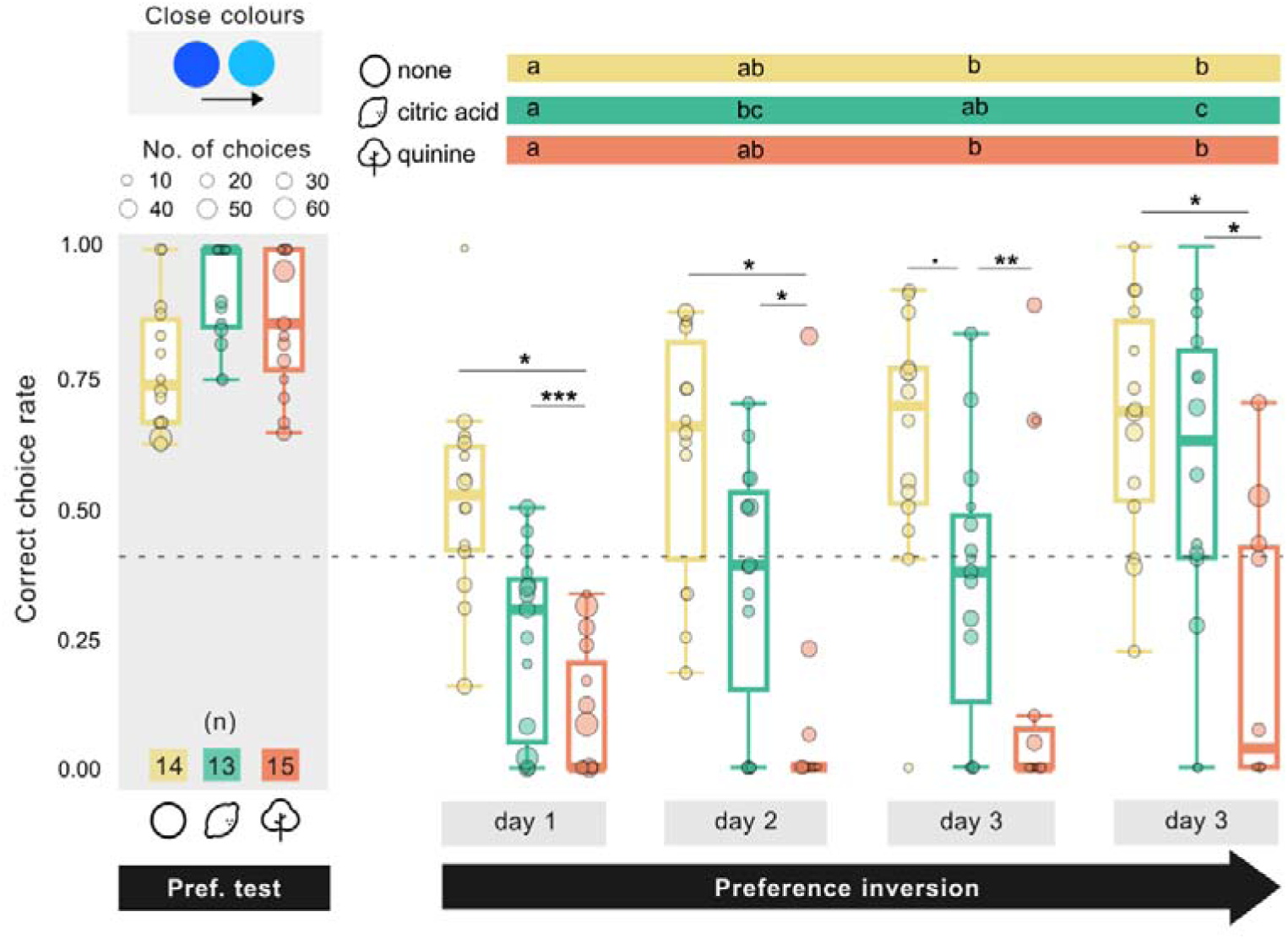
Preference inversion in the close colour experiments. Correct choice rates represent feeding on the preferred colour during the preference test (grey highlight), and on the unpreferred colour (CS+) during the preference inversion phase. Choice rates of individuals are denoted by circles with the size representing the number of choices made in a day (see legend). Statistical results (GLMM, Fig. S4c) of differences in choices rates between days within a treatment type are indicated on the treatment-specific coloured bars where days with no differences (p>0.05) share the same letter. Significant differences between treatment types within a day are shown *with black asterisks* (*** p < 0.001, **< 0.01, *< 0.05,. < 0.1).

Data from the control experiment between close colour stimuli was similarly fit to a binomial GLMM and post-hoc comparisons were performed to estimate choice rate differences between days *(Fig. S2)*. Correlation between the choices on the third and fourth day was calculated with the ‘cor.test’ function using Spearman’s method to assess whether moths made comparable choices in the test with no droplets.

#### Learning curves

To model learning curves, we used a GLMM incorporating a psychometric function (see Kirwan and Nilsson, 2019) using Bayesian estimation. The learning curves were modelled on data from the successful individuals (>40% correct choices on day 4), to visualize how the treatment type affected learning dynamics *(Fig. 4)*. The model was fitted with the Stan language (Carpenter et al., 2017; http://mc-stan.org/) via the brms package (v2.6.0; Bürkner, 2018; https://cran.r-project.org/web/packages/brms/) within R.

**Fig. 4.**
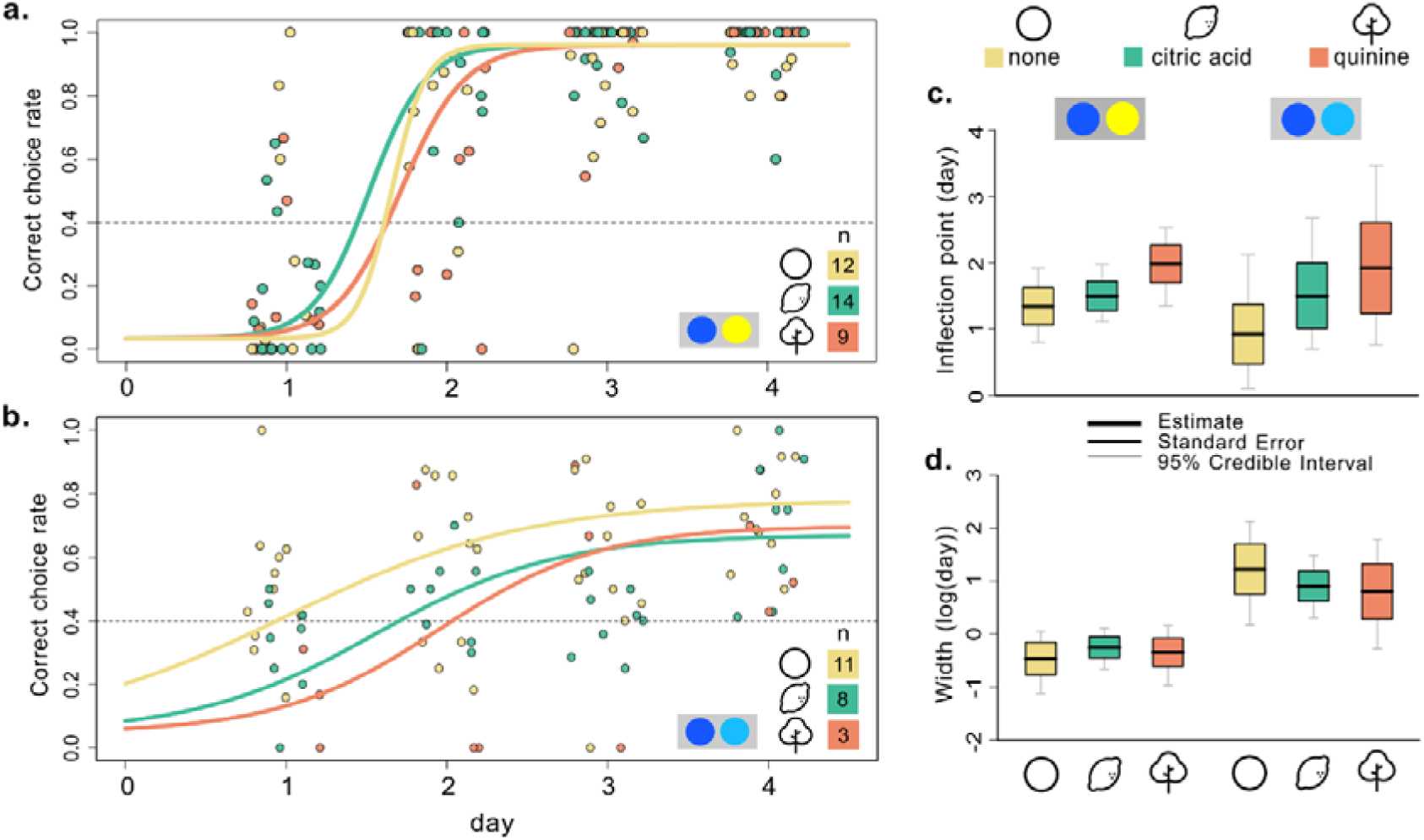
Learning dynamics across treatment types for each discrimination task. **a-b**. Psychometric learning curves modelled for data from successful individuals in the preference inversion phase (see Methods for details). Points depict the correct choice rates of individuals, and colours represent the respective treatment types. Curves are extrapolated to the 0^th^ day to visualize learning predictions for the baseline (initial choice rate). The dotted line depicts the 40% choice criterion which was considered successful learning. **c.** Inflection points of the learning curves estimated for the treatment types for each discrimination task model (close colour and distant colour). Upper and lower box limits depict the Standard error, and the whiskers depict the 95% Credible Intervals. **d.** Widths of the learning curves estimated for the treatment types for each discrimination type model. Blue-yellow and blue-cyan circle pairs depict the discrimination task corresponding to the data.

A logistic regression model modified from Olsson et al. (Olsson et al. 2018) was used, where the initial guess rate and lapse in learning were assigned as the baseline and upper asymptote of the curve respectively. This psychometric formula was used to estimate the learning curve’s inflection point (Inflex) and 80% of the curve’s rise range (Width) to compare between treatment types. To restrict the estimates of the baseline and lapse rate between zero and one, they were estimated on a logit scale {y=ln[x/(1−x)]}, and the curve width was estimated on the natural logarithmic scale. Choices from each day were set as the response variable and fit to a binomial family with an identity link. Informative priors for the baseline were estimated close to 0 based on the animals’ unambiguous choices during the preliminary test. The learning lapse was set near 0 for the distant colour experiments and around 0.25 for the close colour experiments where fewer animals reached a 100% choice rate. Strong priors were also set for the threshold and width of the curve with a distribution centred on the first conditioning day for the distant colour experiments and the third day for the close colour experiments, based on visualising the choice data. Preferred colour, treatment type, choice rate from the preference test, and their interactions were included as fixed effects, and individual ID was set as the random effect to obtain threshold-width values specific to each condition and individual.

A model of best fit was assessed by dropping predictor variables from the full model and comparing all possible models using leave-one-out cross validation via the ‘LOO’ function (Vehtari et al., 2017). The model of best fit was chosen as the one with the highest ELPD (Estimated Log Predictive Density) value. The estimates were calculated from the final model with predictors assigned as

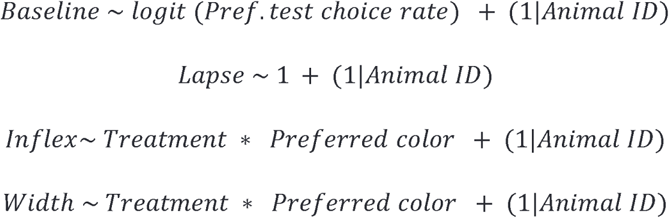

Preferred colour was excluded from the close colour experiments with only blue as the CS-, and based on the data from this discrimination task, the predictors for lapse were adjusted to

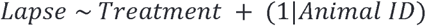

Parameters were estimated using four Markov–Chain Monte Carlo (MCMC) chains running for 2000 iterations post-warmup, and model convergence was checked and adjusted by replacing the default priors with more informative priors when necessary (*Table S2*). The final estimates were drawn from models where all chains converged (R-hat convergence diagnostic < 1.01). Learning curves predicted for each individual were also used to visually validate predictions.

#### Colour switching rates

As a proxy for learning dynamics during the differential conditioning, switching rates were calculated for each individual. Successive choices of the same colour were marked a ‘0’ switch, while choosing different colours in successive flower contacts was marked a ‘1’ switch. The switching rates were then calculated as the sum of switches divided by the number of possible switches across the total choices made by an individual during the differential conditioning period. To examine the variation of switching rates with discrimination and treatment type, a GLM including the treatment types common to both discrimination tasks was modelled using the ‘glmmTMB’ function (Brooks et al., 2017). The model was fit to a betabinomial family with the interaction of treatment and discrimination type, and the number of feeding counts as predictors. After stepwise model reduction, the simplest adequate formula was structured as,

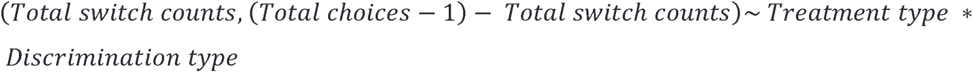

The ‘Anova’ function with Wald’s Chi-square test was used to determine the significance of the fixed variables on the switching rates. Post-hoc comparisons were then conducted to identify pairwise differences using Sidak adjustments for multiple-testing corrections *(Fig. 5b)*.

**Fig. 5.**
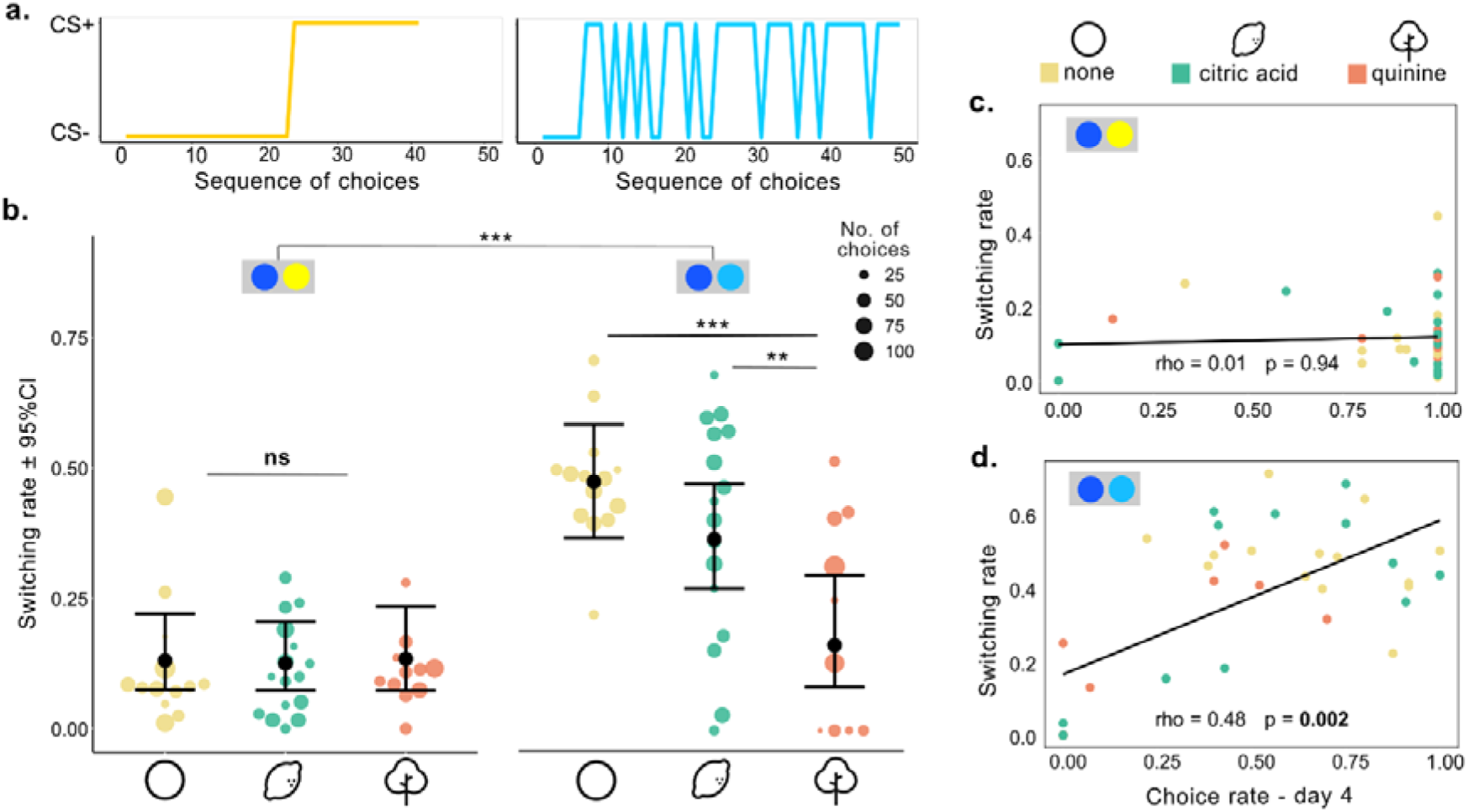
Exploration dynamics during differential conditioning. **a.** Example choice sequence of individual moths in the appetitive-only conditioning from the distant colour (left) and close colour experiment (right). **b.** Switching rates of individuals calculated for choices across the colour inversion phase. A value of 1 represents colour switching with every choice, a value of 0 represents no switching during the entire inversion training. Estimates with 95% confidence intervals (in black) are drawn from the GLMM (see Methods). Switching rate of individual moths are represented as points with the size of the circle depicting the number of choices made by the individual in the colour inversion phase (see legend). *Significant differences* are shown with black asterisks (*** p < 0.001, **< 0.01, *< 0.05). **c-d.** Correlation of switching rates with the choice rate on day 4 of preference inversion. Points represent individual moths, and colours indicate their treatment type. Results of a Spearman’s rank correlation coefficients (rho) are shown in the graph. Blue-yellow and blue-cyan colour pairs depict the discrimination task corresponding to the data.

To test whether switching between colours was linked to the animals’ correct choice rates, we fitted correlations between the switching index and the choices rates on the fourth day using a Spearman’s tests with the ‘cor.test’ function *(Fig. 5c, d)*.

#### Proboscis contact duration

To quantify the duration of proboscis contacts with the treatment substances – citric acid, quinine, water, salt, and the sucrose solution, we annotated videos from the preference inversion phase using BORIS software (v8.22.6, https://www.boris.unito.it/) (Friard & Gamba, 2016). Videos from the first day, during which animals selected both the CS+ and CS-, were collected and scored in a blind assessment where treatment type and the rewarded colour were undisclosed. Contact durations were recorded as time between the first to last moment of contact with a droplet. Only videos where the proboscis contacts were clearly observed were used. To minimise the effects of satiation or habituation on durations, not more than 10 contacts were scored. Only animals with at least 5 contacts with both the CS+ and CS- were included in the final analysis.

A generalised linear mixed model (GLMM) was fitted to a gamma family with a log link with the contact durations as the dependent variable and stimulus type (CS+ or CS-), treatment type, discrimination type, and preferred colour as the predictors, and animal ID as the random effect. Estimates were drawn from the simplest adequate model,

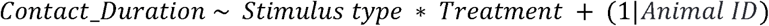

Pairwise comparisons for the treatment types were estimated with Sidak corrections for multiple testing.

## Results

To compare the impact of various negative reinforcements on learning outcome, hawkmoths were trained to invert a colour preference in a discrimination task with either stimuli of perceptually distant colours (blue-yellow) or close colours (blue-cyan). Accordingly, comparisons between the different treatment types, i.e., appetitive-only conditioning, or citric acid, quinine, salt, or water as negative reinforcements were interpreted within their respective discrimination tasks.

Since in all but the appetitive-only conditioning, both stimuli had droplets on the surface, we performed a control with the no substance treatment to ensure that the droplets did not aid associations to the CS+. As there were no significant differences in choice rates between the third day of differential conditioning with only sugar rewards and the subsequent test without droplets on either stimulus (*Fig. S2b*), we concluded that the moths did not rely on the presence of droplets for their choices. A moderate positive correlation (Spearman’s ρ=0.61, p=0.08; *Fig. S2c*) between the choices from the third day and the test supports the conclusion that the animals maintained their associations across days despite the absence of droplets on the flowers. Thus, we conclude that all choices made in these experiments were colour driven.

### Negative reinforcement type did not alter visual discrimination learning of distant colours

In the distant colour experiments, moths pretrained to blue or yellow artificial flowers selected their pretrained colour *(Fig. S3a)* with a high choice rate during the preference test (*Fig. 2 Preference test*).

In the preference inversion phase, moths significantly increased their correct choice rate across the differential conditioning period (GLMM Day - Χ^2^= 570.94, df= 3, p< 0.001; *Fig. 2*). A success criterion was assigned to animals with a 40% baseline of correct choices on day 4, indicating a shift from their initial colour preference of 60%. This criterion was met by 71.05% (54/76) of the conditioned animals with a high median choice rate of 100% *(Fig. 2 Day 4).* The treatment type did not significantly influence the number of conditioned animals that achieved the success criteria (*Fig. S3c, Table S3*).

The moths’ preferred colour influenced their choice rate (GLMM Day: Preferred colour - Χ^2^= 11.62, df= 3, p= 0.009; *Fig. S4b*), with animals conditioned to yellow (CS+) against their blue preference having a greater correct choice rate on the second and third day. This difference disappeared on the last day. Generally, moths with a higher choice rate for their preferred colour during the preference test performed more poorly on the first day of differential conditioning in the inversion phase (Spearman’s test rho= −0.45, p< 0.002; *Fig. S4d*). However, this correlation was not significant by the last day of the conditioning, showing that their initial colour preference did not impact the final learning outcome. We found no interaction between the treatment type and the moths’ preferred colour.

There was also a significant difference in the choices rates across days between treatment types (Day: Treatment - Χ^2^= 40.5465, df= 12, p< 0.001; *Fig. 2*) but within a day there were no significant differences between the treatments. Despite this, we found a general trend of citric acid and appetitive-only conditioned individuals more consistently achieving higher correct choice rates. Quinine appeared to motivate learning more slowly while water and salt shared similar trends with highly variable success rates.

In conclusion, with perceptually distant colours, the negative reinforcement type did not distinctly alter learning success on individual days, as the hawkmoths readily inverted their preferences in all conditions.

### Negative reinforcement type altered visual discrimination learning for close colours

To pose a more challenging learning context, in which the effect of different aversive treatments might be observed more clearly, we conducted the experiment using stimuli closer in the hawkmoths’ colour space (*Fig. S1b*), with treatments that emerged as most effective from the distant colour experiments - citric acid, quinine, and the appetitive-only conditioning.

In the analysis of the close colour experiments, we only included cyan as the CS+, since moths pretrained to both cyan or blue, selected blue flowers during their preference test (*Fig. S3a).* In this discrimination task, fewer moths shifted their colour preference in comparison to the distant colour experiments, with only 52% (22/42) of animals reaching the success criterion. Although the moths modified their foraging choices across the duration of the differential conditioning (GLMM Day - Χ^2^= 47.51, df= 3, p< 0.001; *Fig. 3*), their median choice rate for the CS+ (cyan) by the end of the differential conditioning reached only 51.04% (*Fig. 3 Day 4*), distinctly lower than their counterparts in the distant colour experiments. While successfully conditioned animals in the distant colour experiments decisively chose the CS+, most moths in the close colour experiments did not make as strong an association with the rewarded target.

Interestingly, the treatment type influenced the proportion of animals that reached the success criterion, which was significantly lower when conditioned with quinine in comparison to the appetitive-only condition (Fisher’s exact test: quinine/none- p= 0.01; *Fig. S3b; Table S3*). The treatment type also influenced the variation in choice rates across the differential conditioning period (GLMM Day: Treatment - Χ^2^= 14.862, df= 6, p= 0.021; *Fig. 3*). In contrast to the distant colour experiments, there were significant differences in choice rates between treatment types within each day. Groups with no substance or citric acid on the CS- had a significantly higher correct choice rate on each day than the group treated with quinine (*Fig. 3*). There were no significant differences between the no substance and citric acid groups within days. Since animals in the close colour experiments were trained only with cyan as the CS+, the effect of preferred colour on choice rates could not be assessed.

Thus, with perceptually close colours, where hawkmoths did not learn to invert their colour preference to the same degree as with distant colours, there were distinct effects of the negative reinforcement type on learning: the highest learning success was achieved with the appetitive-only and citric acid treatment, and the lowest with quinine.

### Learning and foraging dynamics during differential conditioning

To investigate the impact of the treatment types on learning dynamics, psychometric learning curves were modelled to the choice data from the successful animals.

The learning curves modelling the distant colour experiments had steep sigmoidal shapes plateauing at a 100% choice rate within the second day of the differential conditioning (*Fig. 4a*). The learning curves of the close colour experiments were flatter and reached their plateau later in the conditioning period at around a 75% choice rate (*Fig. 4b*). Across treatment types, there was a similar trend in the blue-yellow and blue-cyan experiments for the time taken to arrive at the inflection point, the halfway mark between the lower and upper asymptote of the curves. For both discrimination types, colour choices shifted early in the preference inversion phase (*Fig. 4c*), with the inflection point shifted relatively later for quinine, indicating longer conditioning times, while appetitive-only conditioning required the shortest time. This was the case in both colour tasks, but more pronounced with close colours. The widths, which represent the time range during which 80% of changes in the responses occur, were narrow in the distant colour experiments, as represented by the steep learning slopes, and much wider for the close colour experiments where target acquisition occurred at a slower pace (*Fig. 4d*). There were no differences in widths between the treatment types within a discrimination task, suggesting that the treatment type did not significantly change the speed of target acquisition but rather, influenced the time taken by the animals to sample the CS+.

Successful animals in the distant colour experiments rapidly and reliably switched to feeding on the CS+ after a few rewarded encounters. However, in the close colour experiments, most successful individuals distributed their feeding choices between both CS+ and CS-rather than only visiting the rewarded target (*Fig. 5a*). To quantify the likelihood of switching between a preferred and unpreferred colour across both colour discrimination tasks, we calculated the animals’ switching rates, where a value of 1 represented animals that switched colour with every choice, and a value of 0 represented those that exclusively chose one colour during the entire inversion training phase. These measures differed significantly between the treatment types depending on the discrimination tasks within which the data was nested (GLM Treatment type: Discrimination type - Χ^2^= 8.93, df= 2, p= 0.01). The measures indicate a stronger colour fidelity in the distant colour experiment in comparison to the close colour experiments (GLM Discrimination type - Χ^2^= 42.09, df= 1, p< 0.001). Switching rates varied significantly between the treatment types only in the close colour experiments, with appetitive-only conditioning promoting the greatest number of colour switches and quinine the lowest *(Fig. 5b).* We correlated switching rates to task success to assess whether a strong fidelity to their preferred colour could prevent the hawkmoths from sampling the CS+ and forming reward associations. A positive correlation was found between the choice rate on the fourth day of preference inversion and the animal’s switching rate in the close colour experiments (Spearman’s test rho= −0.48, p= 0.002; *Fig. 5d*), but there was no link between switching and success for the distant colour experiments (*Fig. 5c*). Interestingly, we found no correlation between the number of choices made during the conditioning period and the success rates, indicating that their switching rates rather than the number of choices predicted success.

Thus, in addition to learning success, the learning dynamics, both in terms of how fast hawkmoths learned, and how they explored the stimuli, were impacted by the discrimination task and the aversive treatment type. With close colour pairs, appetitive-only conditioning promoted the fastest learning, with the highest number of switches between the two stimuli, while the opposite was true for quinine.

### All treatment types were aversive, but only quinine impacted the positive reinforcement

To quantify whether the moths did, in fact, react aversively upon probing the treatment substances compared to sucrose, and whether there were differences in aversive strengths between the substances, we analysed proboscis contact durations with the droplets across discrimination tasks and treatment types (except appetitive-only, in which we did not present droplets on the CS-). Contact durations were significantly longer for sucrose in comparison to all treatment substances (GLMM Stimulus type - Χ^2^= 226.46, df= 1, p< 0.001; *Fig. 6*) but we found no significant differences in contact durations between the treatment substances. Thus, while the treatment substances were perceived as aversive, there were no treatment-specific differences in feeding responses to indicate disparate aversive strengths. However, the type of negative reinforcement influenced the contacts with sucrose (GLMM Stimulus type: Treatment type - Χ^2^= 29.42, df= 3, p=< 0.001; *Fig. 6*), as quinine-treated animals had significantly shorter feeding durations than animals in all other treatments.

**Fig. 6.**
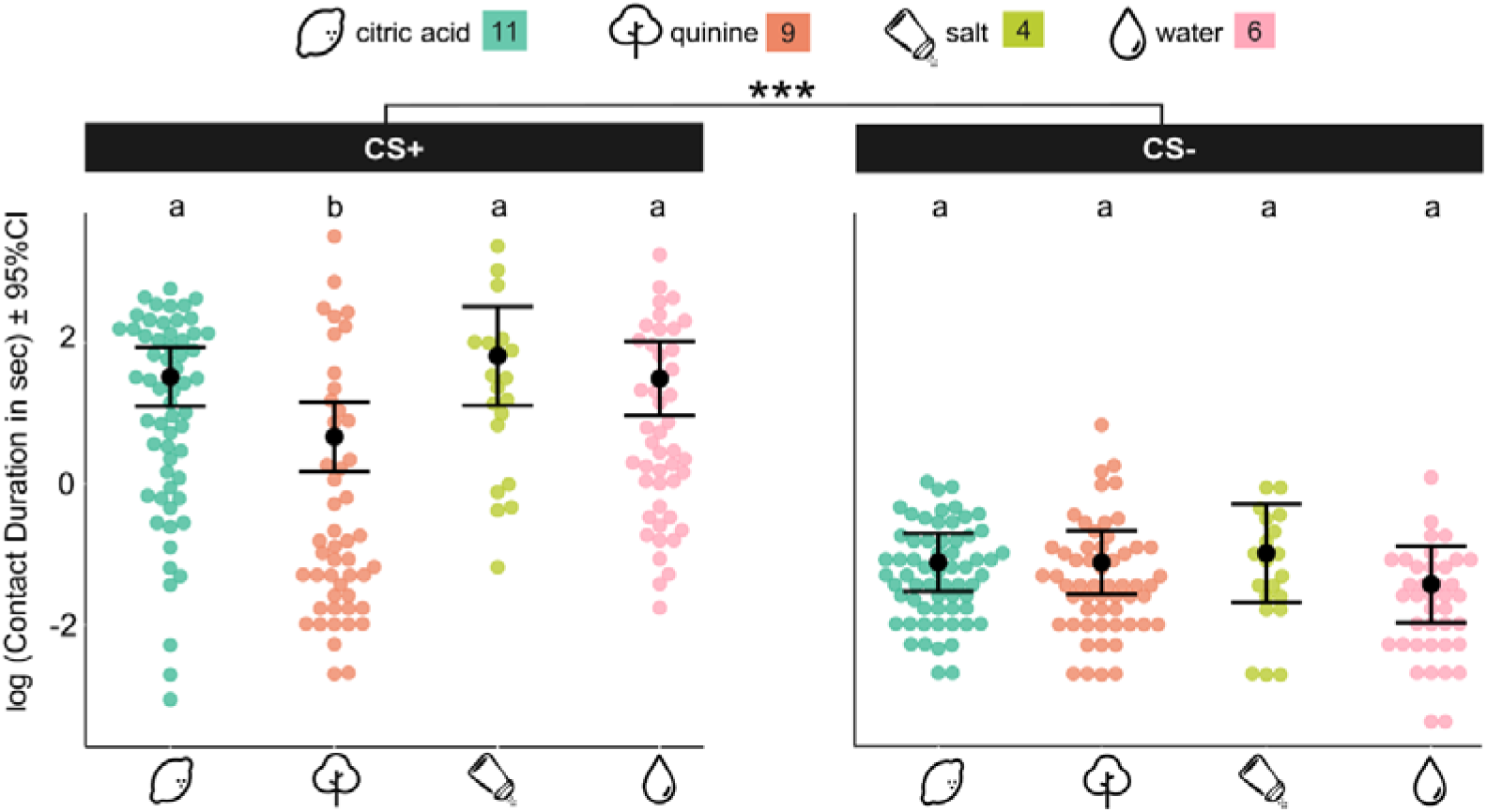
Contact duration with sucrose and the different negative reinforcements. Log transformed contact durations with the droplets on the CS+ (sucrose) and CS- (treatment substances). Shaded points show all the scored contact data for each treatment type. Estimates and their 95% confidence intervals (in black) are drawn from the GLMM (see Methods, Table S6). Differences between treatment types are indicated with letters, where treatments with no difference (p>0.05) share the same letter. Significant difference between the stimulus types is indicated by asterisks (*** p < 0.001).

## Discussion

In this study we demonstrate how the underlying context and conditions can modify learning outcomes by comparing the role of various negative reinforcements for visual discrimination in a solitary insect pollinator. By differentially conditioning the hummingbird hawkmoth, a generalist nectarivore, in two colour discrimination tasks, we show that quinine, a commonly used negative reinforcement, produces worse target association in comparison with other substances, such as citric acid, and with appetitive-only conditioning. We link these differences in learning to the effect of the reinforcing substance on modifying reward association and foraging dynamics in freely flying animals, subsequently affecting the individual’s accuracy and speed of learning.

### Learning asymmetry and innate preferences

Learning dynamics between the close and distant colour experiments varied greatly, with animals making rapid and reliable associations to the rewarded blue or yellow but failing to make similar associations to the cyan flowers. These results correspond to previously reported colour discrimination results in honeybees and bumblebees (Avarguès-Weber et al., 2010; Dyer & Chittka, 2004) where discrimination thresholds for perceptually close colours do not reach the same criteria as those of distinct colours, even with strongly reinforced training protocols.

In our experiments, the differences between the colour discrimination tasks are unlikely to be explained by the challenge of perceptually distinguishing the colours, as *M. stellatarum* can distinguish spectrally similar colours differing by as low as 1-2 nm in the blue 400 and 480 nm range, a spectral resolution even higher than in honeybees (Telles et al., 2016), and can also use achromatic cues to discriminate stimuli (Kelber, 2005; van der Kooi & Kelber, 2022). The cyan and blue stimuli we used had both achromatic and chromatic differences that were large enough to be exploited by the hawkmoths, and correspondingly, they made strongly biased colour choices during their initial preference test, which would not be possible if they were unable to discriminate between the colours.

Rather, the differences in choice rates between the distant and close experiments could have emerged from the moths’ strong innate colour bias, since individuals show untrained preferences for blue or yellow (Kelber, 1996, 2002). The cyan stimulus used in this setup was less preferred in comparison, as evidenced by the results of the pretraining where even after being absolutely conditioned to cyan for 3 days, moths chose to feed on blue when given a choice. While the change from their initial choice rates suggests that learning did take place in both discrimination tasks, new reward associations with the cyan flowers might have been created without losing their initial preference for blue. As a result, at the end of the differential conditioning period, most moths chose equally between cyan and blue rather than associating with a single colour as with the blue-yellow stimuli. Innate colour preferences could have also led to the asymmetric learning acquisitions in the distant colour experiments between the groups with blue and yellow as their CS+. In this species, a larger proportion of animals have a spontaneous preference for blue over yellow (Kelber 1997). Subsequently, more moths with an innate blue preference would have been pretrained through absolute conditioning to yellow artificial flowers than the other way around. Thus more animals with blue as their CS+ would have been ‘re-learning’ a colour, a task requiring more cognitive flexibility and considered more challenging (Izquierdo et al., 2017; Raine & Chittka, 2012; Young et al., 2024).

Another possible explanation for the decreased target acquisition in the close colour experiments could emerge from the moths allocating their attention to the salient blue stimulus and not forming immediate associations to the rewarded target. Such phenomena as blocking or overshadowing of learning due to the presence of a stimulus with high valence in the same environment have been suggested in previous learning studies (Balkenius & Kelber, 2006) in this species.

An interesting aspect of the hawkmoths’ colour associations is demonstrated by their foraging dynamics, where we found a low colour fidelity in the close colour setup, whereas animals rarely switched between the yellow and blue colours. This could indicate a categorical grouping of colours, analogous to human-defined groups, where colours across a range of chromatic and achromatic properties are grouped into the same class, such as blues, greens, yellows, etc.. Seen through the lens of foraging, this would suggest that hawkmoths are less prone to switching between flowers in different colour categories, but when their preferred colour is unavailable or unrewarded, might sample instead from flowers with similar colours. While categorical colour perception has been uniquely linked to primates (Kelber, 2016), generalization of colours by insects, where behavioural responses do not correlate to the colours’ distance in their colour space (Benard & Giurfa, 2008; Troje, 1993), might show that similar processes are utilized.

Tasks requiring discrimination between close colours have been described as ‘challenging’ due to the lower learning or target acquisition levels and have subsequently been used to demonstrate the benefit of strong negative reinforcements for learning in insects (Avarguès-Weber et al., 2010; Dyer & Chittka, 2004; Dyer & Howard, 2023; Giurfa, 2004). However, what is viewed as a learning deficit might instead be the animal’s strategy to benefit from exploring within a colour category rather than making strong associations to a particular colour within a category and restricting foraging flexibility. Here, we interpret the results of discrimination learning as an interaction of their innate preferences, which are known to be reliably recalled even after training or hibernation periods (Gumbert, 2000; Kelber, 2010b; Menzel, 1968), and the functional basis of colour recognition for hawkmoths while foraging.

### Tastes as negative reinforcements in hawkmoths

Colour discrimination learning was influenced by the type of negative reinforcement used during the differential conditioning, which points to a disparate negative valence associated with the treatment substances. The 5-way comparison of treatments in the distant colour experiment, albeit statistically inconclusive, produced citric acid, quinine, and appetitive-only conditioning as the more reliable training conditions, suggesting that these substances were perceived differently from at least water and salt solution. The next series of experiments, where moths were trained to discriminate close colours, demonstrate that citric acid and quinine were also perceived differently from each other. This points strongly to aversion being generated not only due to the mismatch of the substance to the expectation of a sugar reward, but also due to the specific substances being perceived uniquely. Such differences in perception could emerge from a negative sensory experience on sampling the substance, such as that caused by an aversive taste or viscosity (Josens & Farina, 2001). The hawkmoth’s proboscis have sensilla with gustatory and mechanosensory receptors which they use to assess food quality (Kelber, 2003). Studies on the gustatory coding of a closely related hawkmoth, *Manduca sexta*, suggest that tastes in this species are represented in a tastant-dependent manner rather than in the categories - bitter, salt, sour, sugar, umami-typically known in other animals (Reiter et al., 2015). In another moth, *Heliothis virescens*, it was shown that two bitter substances, one of which was quinine, produced separate behavioural and neural responses (Kvello et al., 2010). We did not find differences in the proboscis contact durations between the treatment substances which could have been due to the limitation of visually scoring feeding from the videos, where finer differences in contact durations might not have been captured. It is also possible that all substances were aversive beyond a threshold at which the moths show highly stereotyped behaviour of quickly retracting their proboscis. However, the differences in learning outcomes as well as previous studies on moth gustation suggest that the negative reinforcements were perceived as unique gustatory sensations.

Post-ingestional physiological effects, such as malaise, or a degradation of the reward quality of sugar from contacting the treatment substances could have also played a role in learning. In our experiments, contact durations with the treatment substances were significantly lower than those with the sucrose rewards and typically lasted less than a second, making it unlikely that they ingested toxic quantities of the substance. Instead, upon contacting quinine on the CS-, moths made much shorter contacts with sucrose and were observed to sometimes be repelled on contacting the sucrose droplets. This strongly evidences the inhibition of sugar receptors by bitter substances that has been reported in many insects (De Brito Sanchez et al., 2005; Dethier & Bowdan, 1992; Jørgensen et al., 2007; Liscia & Solari, 2000; Meunier et al., 2003). Thus, using quinine as a negative reinforcement could have prevented animals from making accurate associations to the rewarded stimulus. Additionally, we also found a lower switching rate for quinine reinforced animals with the close colour stimuli which could have been due to the lack of motivation to diverge from their preferred colour since the alternate was also not perceived as rewarding.

### Learning visual discrimination

Previous work on insects suggests that differential conditioning using strongly aversive substances improves target recognition (Avarguès-Weber et al., 2010; Dyer & Chittka, 2004; Dyer & Howard, 2023; Rodríguez-Gironés et al., 2013) as a consequence of the punishment associated with making incorrect choices, along with the positive conditioning to the rewarded reinforcement. The faster and stronger associations thus formed are discussed as emerging from the punished individual being more attentive to the task (Chittka et al., 2003) and retaining more information associated with the stimuli (Giurfa, 2004). Although learning studies in Lepidoptera have demonstrated that feeding behaviour can be modified in response to non-sugar compounds (Broadhead & Raguso, 2021; Rodrigues et al., 2010), in our experiments, moths treated with strong negative reinforcements such as concentrated quinine or salt solution did not perform better than the animals that only encountered an empty flower. Rather, finding their preferred colour sugar-empty motivated the animals to sample the alternative colour, resulting in better colour associations with the rewarded target. Interestingly, we found no correlation between the number of foraging attempts and the learning acquisition, but rather, in the present discrimination context, the animal’s tendency to explore foraging alternatives and encounter the rewarded target predicted task success

Hawkmoths are solitary insects making few and therefore more restricted feeding choices in comparison to many eusocial foragers that make multiple foraging bouts to feed the colony. Moreover, insects within a eusocial lifestyle may display more exploratory behaviour since they may also supplement their nutrition and fitness through the hive (Hendriksma et al., 2019), whereas risk aversion could be a strong limiting factor in the foraging behaviour of solitary pollinators. In the case of the hummingbird hawkmoth, we found that conditioning with an aversive stimulus diminished their motivation to explore foraging alternatives in the vicinity of the stimulus, instead of providing association cues to aid visual discrimination. Since hawkmoths form feeding associations in a highly reward-motivated manner (Balkenius et al., 2008; Kelber, 1996), with the ability to discriminate between sugar types and learn colours associated with their preferred sugars (Kelber, 2003), training protocols with disparate reward associations might motivate learning, while also preventing any loss in performance caused by harsh punishments. Such a conditioning method would also represent a naturalistic foraging situation for this species, as their longer lifespans and migration activities (Kelber, 2010b) require them to adapt to new environments with different resources and select among the most energy efficient food source.

The contrast in learning outcomes between eusocial and solitary pollinator species in response to negative reinforcements suggests that aversion does not pervasively enhance associations across insect groups. This highlights the necessity of interpreting learning and foraging behaviour not only through the animals’ sensory ecology, but also as a factor of their life history.

### Conclusions

This study highlights that visual learning in insects can be strongly influenced by the underlying context and conditions. Contrary to findings in eusocial insects where learning success has been linked to differential conditioning with strong negative reinforcements, hummingbird hawkmoths showed poorer target association when trained with quinine, compared to citric acid or appetitive-only conditioning with no aversive substance. In addition to interfering with sucrose perception and thus hindering reward association, we demonstrate that quinine was not conducive to the hawkmoths’ exploratory behaviour during foraging. These results underscore the role of foraging dynamics and an animal’s sensory ecology in shaping learning outcomes.

## Supporting information

https://figshare.com/s/4015a0896e8a9366ed72

## Acknowledgements

We thank Sophie Fetter for assistance with scoring videos, and Samira Giger and Lucas Heger for helping with pilot studies to establish methods. We are also grateful to Valerie Kuklovsky for providing useful input, and to all the students involved with animal care during this study.

## Author contributions

Conceptualization: A.N.M., A.S.; Methodology: A.N.M., A.S., J.J.F.; Validation: A.N.M., A.S.; Formal analysis: A.N.M, J.J.F.; Investigation: A.N.M.; Resources: A.S.; Data curation: A.N.M.; Writing - original draft: A.N.M.; Writing - review & editing: A.N.M., A.S., J.J.F.; Visualization: A.N.M.; Supervision: A.S.; Funding acquisition: A.S

## Supplementary Material

Figures S1 to S4. Tables S1 to S6.

