## Supplementary material for "Strong negative reinforcement interferes with visual learning in a solitary pollinator": https://figshare.com/s/4015a0896e8a9366ed72

Supplementary figures

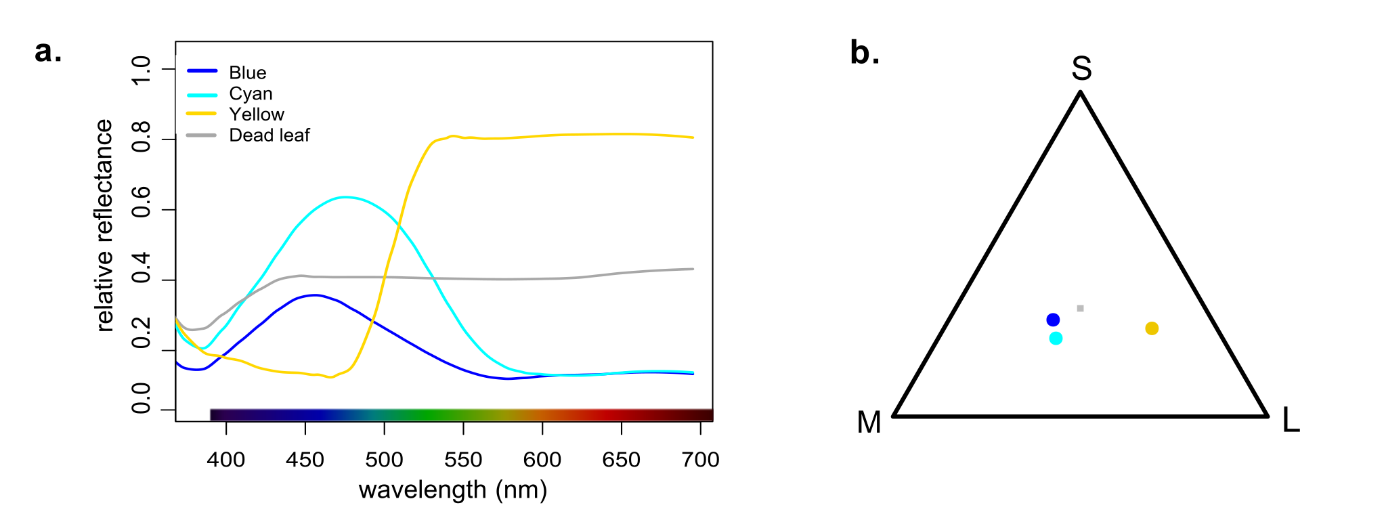

###### Fig. S1 **Spectral characteristics of the stimuli.** **a.** Relative radiance of the stimuli and the dead-leaf background. **b**. Relative stimulation by the blue, cyan, and yellow stimuli (shown as coloured points) to the UV (S), blue (M), and green (L) sensitive photoreceptors in hummingbird hawkmoths (for details on colour vision modelling, see Methods).

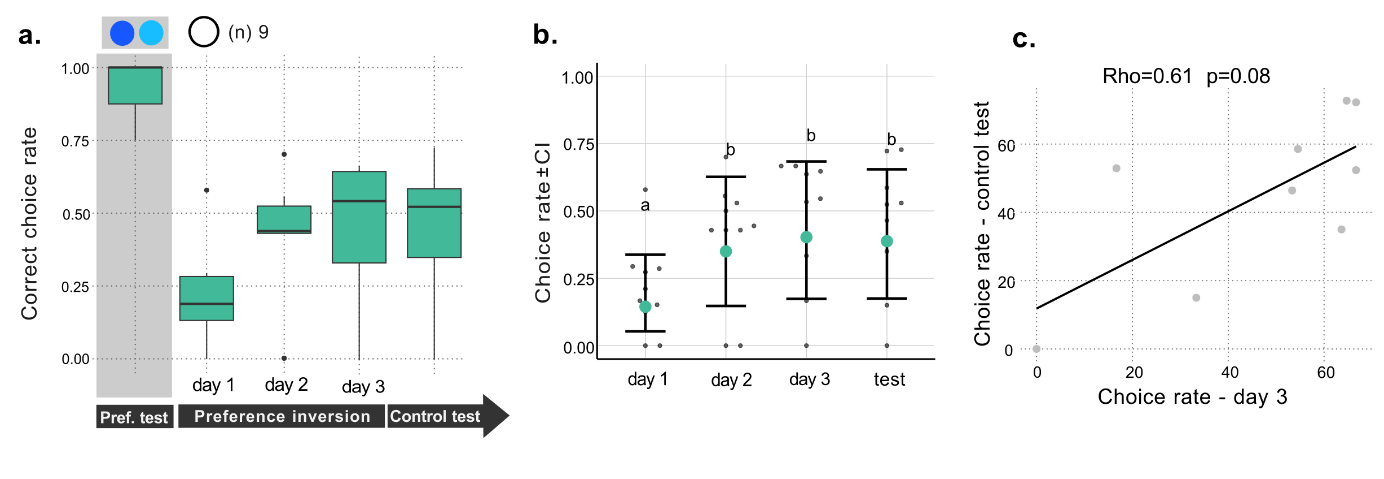
Fig. S2 **Control experiments with the appetitive-only condition** **a.** Choice rates across the preference test, inversion phase, and unrewarded test. Correct choices are marked as feeding on the preferred colour during the preference test, and on the unpreferred colour (CS+) during the preference inversion phase. **b.** Estimated choice rates with 95% confidence intervals (in black) from the GLMM (see Methods). Choice rates of individual moths across days are represented as black dots. **c.** Correlation of choice rates during the unrewarded test and the choice rate on day 3 of the preference inversion. Gray dots represent data from individual moths.

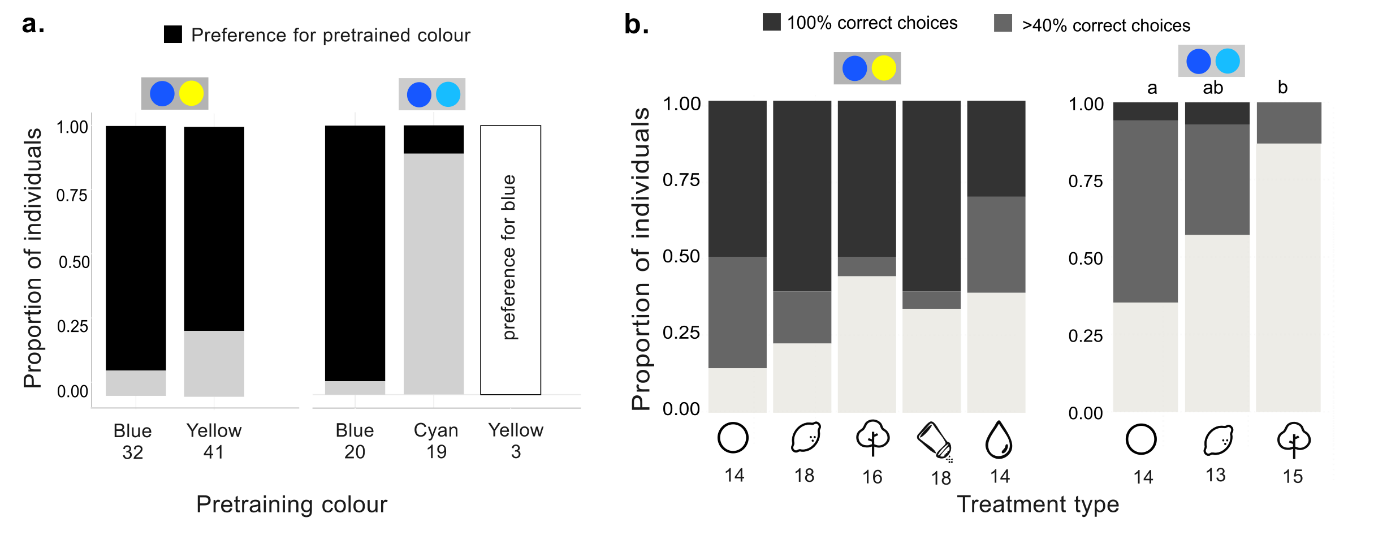

###### Fig. S3 **Successful animals from the pretraining and preference inversion phase.** **a**. Proportion of animals that chose the same stimulus colour as the one they were pretrained to. Sample sizes for the groups are given below. 3 animals that were pretrained using yellow were tested for a preferred colour between blue and cyan and transferred to the close colour experiments. **b**. Proportion of differentially conditioned individuals achieving the success criteria on day 4 (>40% correct choices in grey, and 100% correct choices in black). Pairwise comparisons (Fisher’s test) of individuals with the 40% choice rate found no significant differences between treatment types in the distant colour experiments. Significant differences (p>0.05) in close colour experiments are represented as letters. Blue-yellow and blue-cyan colour pairs depict the discrimination type of the data.

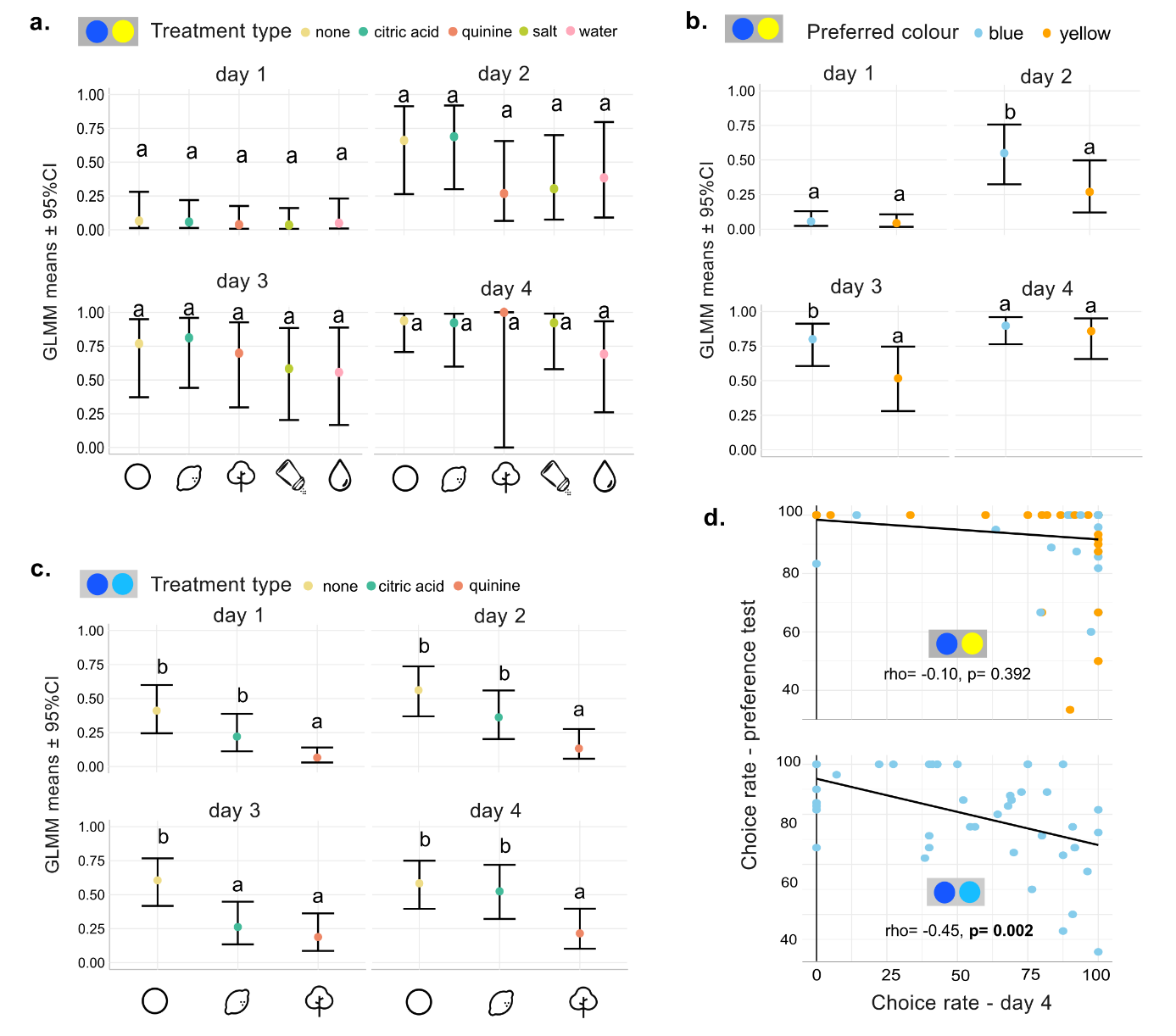

###### Fig. S4 **Model results of the distant and close colour discrimination tasks** **a-c** Estimated choice rates from the GLMM with 95% confidence intervals (in black). Significant differences between treatment types (a and c), or between preferred colours (b) within a day are indicated with letters. Groups with no differences (p>0.05) share the same letter. **d.** Correlation between the choice rates for the preferred colour during the preference test and for the CS+ on day 4 of the preference inversion phase. Results of a Spearman’s rank correlation coefficients (rho) are shown above the graph. Blue-yellow and blue-cyan colour pairs depict the discrimination type of the data. distant and close, respectively. Animals with blue as their preferred colour (CS-) are represented by blue dots, while animals with yellow as their preferred colour are represented by orange dots.

### Supplementary tables

###### Table S1 Relative chromatic and achromatic distances between the stimuli calculated as noise-weighted chromatic Euclidean distances measured as Just Noticeable Differences (JNDs) in the hawkmoth colour space.

| Colour patch 1 | Colour patch 2 | Chromatic distance | Achromatic  distance |
| --- | --- | --- | --- |
| Blue | Cyan | 1.18 | 6.68 |
| Blue | Yellow | 5.99 | 10.13 |
| Cyan | Yellow | 5.42 | 3.45 |

###### Table S2 Priors assigned to model psychometric learning curves for the colour inversion phase. Informative priors were assigned to the intercepts and standard deviations of the Inflex, Width, Base and Lapse rates of the curves based on choice data from the respective discrimination tasks: Close and Distant colours.

| Discrimination type | prior | class | coef (Group) | nlpar |
| --- | --- | --- | --- | --- |
| Close colors | student_t(3, 0, 1.0) | b |  | Inflex |
|  | normal(2,0.8) | b | Intercept | Inflex |
|  | normal(0,1) | b | Treatmentnone | Inflex |
|  | normal(0,1) | b | Treatmentquinine | Inflex |
|  | student_t(3, 0, 1.0) | sd |  | Inflex |
|  | student_t(3, 0, 1.0) | sd | Individual | Inflex |
|  | student_t(3, 0, 1.0) | sd | Intercept Individual | Inflex |
|  | normal(0,1) | b |  | logitBase |
|  | normal(-2.5,1) | b | Intercept | logitBase |
|  | normal(0,0.5) | b | logitPrelim | logitBase |
|  | student_t(3, 0, 0.5) | sd |  | logitBase |
|  | student_t(3, 0, 0.5) | sd | Individual | logitBase |
|  | student_t(3, 0, 1.0) | sd | Intercept Individual | logitBase |
|  | normal(0,1) | b |  | logitLapse |
|  | normal(-1,0.5) | b | Intercept | logitLapse |
|  | normal(0,1) | b | Treatmentnone | logitLapse |
|  | normal(0,1) | b | Treatmentquinine | logitLapse |
|  | student_t(3, 0, 1) | sd |  | logitLapse |
|  | student_t(3, 0, 1) | sd | Individual | logitLapse |
|  | normal(0,1) | sd | Intercept Individual | logitLapse |
|  | normal(0,1) | b |  | logWidth |
|  | normal(log(2),0.3) | b | Intercept | logWidth |
|  | normal(0,1) | b | Treatmentnone | logWidth |
|  | normal(0,1) | b | Treatmentquinine | logWidth |
|  | student_t(3, 0, 1.0) | sd |  | logWidth |
|  | student_t(3, 0, 1.0) | sd | Individual | logWidth |
|  | student_t(3, 0, 1.0) | sd | Intercept Individual | logWidth |
| Distant colors | student_t(3, 0, 1.0) | b |  | Inflex |
|  | normal(1.5,1.0) | b | Intercept | Inflex |
|  | normal(0,0.5) | b | Preferred_colory | Inflex |
|  | normal(0,0.5) | b | Treatmentnone | Inflex |
|  | normal(0,0.5) | b | Treatmentnone:Preferred_colory | Inflex |
|  | normal(0,0.5) | b | Treatmentquinine | Inflex |
|  | normal(0,0.5) | b | Treatmentquinine:Preferred_colory | Inflex |
|  | normal(0,0.5) | b | Treatmentsalt | Inflex |
|  | normal(0,0.5) | b | Treatmentsalt:Preferred_colory | Inflex |
|  | normal(0,0.5) | b | Treatmentwater | Inflex |
|  | normal(0,0.5) | b | Treatmentwater:Preferred_colory | Inflex |
|  | student_t(3, 0, 1.0) | sd |  | Inflex |
|  | student_t(3, 0, 1.0) | sd | Individual | Inflex |
|  | normal(0,0.5) | sd | Intercept Individual | Inflex |
|  | normal(0,0.5) | b |  | logitBase |
|  | normal(-2.5,1.0) | b | Intercept | logitBase |
|  | normal(0,0.2) | b | logitPrelim | logitBase |
|  | student_t(3, 0, 4.4) | sd |  | logitBase |
|  | normal(0,0.5) | sd | Individual | logitBase |
|  | student_t(3, 0, 0.5) | sd | Intercept Individual | logitBase |
|  | normal(0,0.5) | b |  | logitLapse |
|  | normal(-4,1.0) | b | Intercept | logitLapse |
|  | student_t(3, 0, 1.0) | sd |  | logitLapse |
|  | student_t(3, 0, 1.0) | sd | Individual | logitLapse |
|  | student_t(3, 1.0, 1.0) | sd | Intercept Individual | logitLapse |
|  | normal(0,0.5) | b |  | logWidth |
|  | normal(log(1.0),1.0) | b | Intercept | logWidth |
|  | normal(0,0.5) | b | Preferred_colory | logWidth |
|  | normal(0,0.5) | b | Treatmentnone | logWidth |
|  | normal(0,0.5) | b | Treatmentnone:Preferred_colory | logWidth |
|  | normal(0,0.5) | b | Treatmentquinine | logWidth |
|  | normal(0,0.5) | b | Treatmentquinine:Preferred_colory | logWidth |
|  | normal(0,0.5) | b | Treatmentsalt | logWidth |
|  | normal(0,0.5) | b | Treatmentsalt:Preferred_colory | logWidth |
|  | normal(0,0.5) | b | Treatmentwater | logWidth |
|  | normal(0,0.5) | b | Treatmentwater:Preferred_colory | logWidth |
|  | student_t(3, 0, 1.0) | sd |  | logWidth |
|  | student_t(3, 0, 1.0) | sd | Individual | logWidth |
|  | student_t(3, 0, 1.0) | sd | Intercept Individal | logWidth |

###### Table S3 Fischer’s exact test for pairwise comparisons between treatment types (Group 1 and 2) to compare number of successful (>40% choice rate on day 4) and unsuccessful individuals in the differential conditioning. Significant p-values are highlighted.

| Discrimination type | Group1 | Group2 | OddsRatio | CI_Lower | CI_Upper | P_Value | P_Adjusted |
| --- | --- | --- | --- | --- | --- | --- | --- |
| Distant colours | citric acid | quinine | 2.64 | 0.27 | 34.66 | 0.27 | 1 |
|  | citric acid | salt | 1.57 | 0.13 | 21.42 | 0.70 | 1 |
|  | citric acid | water | 2.13 | 0.17 | 31.06 | 0.43 | 1 |
|  | citric acid | none | 0.59 | 0.01 | 11.05 | 0.67 | 1 |
|  | quinine | salt | 0.59 | 0.05 | 5.70 | 0.71 | 1 |
|  | quinine | water | 0.81 | 0.07 | 8.53 | 1 | 1 |
|  | quinine | none | 0.23 | 0.00 | 3.03 | 0.12 | 1 |
|  | salt | water | 1.36 | 0.11 | 16.87 | 0.71 | 1 |
|  | salt | none | 0.38 | 0.01 | 6.01 | 0.40 | 1 |
|  | water | none | 0.28 | 0.01 | 4.65 | 0.21 | 1 |
| Close colours | citric acid | quinine | 5.93 | 0.67 | 80.00 | 0.05 | 0.10 |
|  | citric acid | none | 0.45 | 0.03 | 4.50 | 0.42 | 0.42 |
|  | **quinine** | **none** | 0.08 | 0.00 | 0.72 | 0.00 | **0.01** |

###### Table S4 Generalised Linear Mixed Model (GLMM) for the choice rates from the differential conditioning experiments. Analysis of deviance is reported with type 2 Wald ‘s chi-square tests. Significant p-values are highlighted.

| Discrimination type | Parameters | Χ^2^ | Df | p-value |
| --- | --- | --- | --- | --- |
| Distant colours | **Day** | 670.76 | 3 | **< 2.2e-16** |
|  | Treatment | 3.82 | 4 | 0.60 |
|  | Preferred_colour | 2.51 | 1 | 0.18 |
|  | Preference test_choice rate | 3.65 | 1 | 0.10 |
|  | **Day:Treatment** | 40.55 | 12 | **0.002** |
|  | **Day:Preferred colour** | 9.07 | 3 | **0.002** |
| **Close colours** | **Day** | 48.84 | 3 | **2.026e-10** |
|  | **Treatment** | 23.33 | 2 | **8.572e-06** |
|  | **Preference test_choice rate** | 6.43 | 1 | **0.01** |
|  | **Day:Treatment** | 14.96 | 6 | **0.02** |

###### Table S5 Generalised Linear Model (GLM) for the switching index from the differential conditioning experiments in the close and distant colour experiments (Discrimination type). Analysis of deviance is reported with type 2 Wald ‘s chi-square tests. Significant p-values are highlighted.

| Parameters | Χ^2^ | Df | p-value |
| --- | --- | --- | --- |
| **Treatment** | 8.34 | 2 | **0.015** |
| **Discrimination type** | 41.75 | 1 | **1.04e-10** |
| **Treatment:Discrimination type** | 8.89 | 2 | **0.011** |

###### Table S6 Generalised linear Mixed Model (LMM) for the contact durations on the stimulus (CS+ or CS-) from the differential conditioning experiments. Analysis of deviance is reported with type 2 Wald ‘s chi-square tests. Significant p-values are highlighted.

| Parameters | Χ^2^ | Df | p-value |
| --- | --- | --- | --- |
| **Stimulus** | 285.56 | 1 | **< 2.2e-16** |
| **Treatment** | 14.48 | 3 | 0.163 |
| **Stimulus:Treatment** | 29.42 | 1 | **0.00034** |
